# Developing Novel Biologicals with Multiple Modes of Action (BM) to Combat the Early Blight Disease in Tomato (*Solanum lycopersicum*), Potato (*Solanum tuberosum*), and Bell Pepper (*Capsicum annuum*)

**DOI:** 10.1101/2022.10.29.514359

**Authors:** Shloke Patel, Balaji Aglave

## Abstract

Early blight, a fungal disease, leads to annual crop losses of up to 79% in tomato, potato, and bell pepper crops. Current treatment requires repeated application of chemical fungicides, resulting in fungicidal resistance. Previous studies have identified microorganisms that enhance plant growth or prevent the growth of early blight spores. This study aims to develop a biological fungicide with multiple modes of action (BM) that can be used in fungicide rotation. The average area under the disease progress curve (AUDPC) values were calculated to quantify the disease progression; formulation 2, formulation 3, and formulation 3 produced the lowest AUDPC value for the tomato, potato, and bell pepper crops, respectively. All the treatments were compared with the current industry standard used to treat early blight, and analysis of variance (ANOVA) tests, with 95% confidence intervals (CIs), of the disease severity were conducted to determine the efficacy of the treatments.

## 1. Introduction

Tomato, potato, and bell pepper are some of the most economically important crops globally. As members of the nightshade family of plants, they are susceptible to infection by *Alternaria solani,* the fungal pathogen that causes early blight (Bauske et al., 2018). Economic yield losses due to early blight have been recorded at 79% in some cases (Adhikari et al., 2017).

Early blight is a foliar disease primarily afflicting plants grown in humid climates and sandy soil. Temperatures from 23.89°C to 28.89°C are optimal for the development of early blight (Li, 2012). The fungus is spread through contaminated soil and plant debris, and the most common symptoms of early blight are the formation of dark, concentric lesions on the plant’s foliage and stems from which spores are produced and spread to other crops by wind or irrigation (Stevenson et al., 2002). Current treatment for plant-parasitic fungi is intensive, involving frequent chemical fungicide applications and heavy monitoring of the crop foliage and roots. The treatment and prevention of early blight require copious amounts of chemical fungicides to be regularly sprayed in the fields, increasing the likelihood of the development of fungicidal resistance (Hahn, 2014).

Farmers have tried to control the spread of this fungus by using chemical fungicides such as Endura® and Inspire Super^®^. However, the sequential use of these potent fungicides is limited in the United States by the Environmental Protection Agency (EPA) because of their hazardous effects on the environment and can result in nutrient depletion in the soil and contamination of surrounding water bodies because of their active ingredients (Zubrod et al., 2019). Endura^®^ contains boscalid, and Inspire Super^®^ contains difenoconazole (Kish, 2008; Garvie, 2019). Microbial formulations can be a potential organic alternative to effectively control the early blight fungus and limit the environmental damage of the fungicidal treatments (Zubrod et al., 2019), and in this investigation, three novel microbial formulations were developed (Table 1) and tested against early blight development on tomato, bell pepper, and potato. The advantage of microbial formulations over chemical fungicides is the multiple modes of action provided by the microbial cocktails (Köhl, 2019).

**Table 1.** Summary of the treatments used in this study.

|  |  |
| --- | --- |
| Formulation 1 | *Bacillus subtilis* 29784, *Bacillus subtilis* 168, *Azotobacter* sp. |
| Formulation 2 | *Glomus claroideum*, *Glomus etunicatum*, *Bacillus subtilis* 29784 |
| Formulation 3 | *Trichoderma viride*, *Glomus viscosum*, *Acetobacter* sp. |
| Industry Standards | Inspire Super® (difenoconazole) and Endura® (boscalid) |
| Untreated Control | Sterile distilled water |

## 2. Materials and Methods

### 2.1 Fungal cultivation

*Glomus claroideum, Glomus etunicatum, Trichoderma viride,* and *Glomus viscosum* were extracted from soil, in the field at Florida Ag Research, using the serial dilution technique (Ben-David and Davidson, 2014) and individually cultivated on potato dextrose agar (PDA) plates using two mycelia discs per plate (twenty plates total for each strain). The inoculated plates were incubated in a laminar flow biosafety hood at 37 °C for 2 weeks. The mycelia on the agar plates were harvested by excision with a sterile scalpel. The mycelia from each strain were placed separately in a flask containing 100 mL of sterile distilled water. The total number of spores in each suspension was counted and adjusted to 3 × 10^8^ per mL. The flasks were placed in a shaker incubator at 1000 rpm for 6 hours at an interval of 30 minutes on and 30 minutes off daily for 1 week to cultivate the fungal strains. The microbial formulations were prepared by mixing each strain according to the compositions in Table 1. The early blight cultivation procedure followed the same method as the fungal cultivation procedure described above with the inclusion of an additional cultivation on 400 PDA plates prior to spore collection and enumeration.

### 2.9 Bacterial cultivation

For the bacterial cultivation and formulation development, 20 petri plates prepared with PDA were used for each microorganism and were inoculated with 2 bacterial discs which were pure cultures (Table 1) from the soil using the serial dilution technique and validated by the gram staining technique (Tripathi, 2021); only the *Bacillus subtilis 29784* and *Bacillus subtilis 168* bacteria were sourced from the ATCC Bank and were validated by the acid-fast staining technique. For the initial replication, the microorganisms were stored in a laminar air flow at 37 °C for 2 weeks. After 2 weeks, 320 colonies were collected into 2 L of nutrient broth and were stored in the laminar flow for 2 weeks. The 2 L of the nutrient broth, for each microorganism, was divided into 8 250 mL conical flasks, and the total number of spores were counted to 2 × 10^8^ per mL and the flasks were placed in a shaker incubator at 37°C for 72 hours.

### 2.9 Formulation Preparation

The microorganisms were mixed according to the formulation compositions (Table 1) and were diluted by a 1:4 ratio of microbial formulation to distilled sterile water. The formulations were stored in the refrigeration at 4 °C.

### 2.4 Enumeration of Fungal Spores and Bacterial Cells

To assure that the spore and bacterial concentration of the formulations was constant throughout this investigation, a hemocytometer was used to assess and alter the bacterial and fungal formulations. Initially, 100 μL of the formulation, bacterial or fungal, was placed in a microcentrifuge tube and the formulation was diluted with 100 μL of trypan blue to differentiate dead and living spores or bacterial cells. Then pipette the mixture from the microcentrifuge tube under the cover slip, into the cells of the hemocytometer, the spores or bacterial cells were counted from the 4 corners as well as the center, shown in Figure 1, the cells that were touching the boundaries of the squares were ignored. The concentration was calculated using the formulas displayed in Table 2, the average number of cells per square was calculated, then the dilution factor was calculated, and those values were inputted into the formula to calculate the concentration.

**Figure 1.**
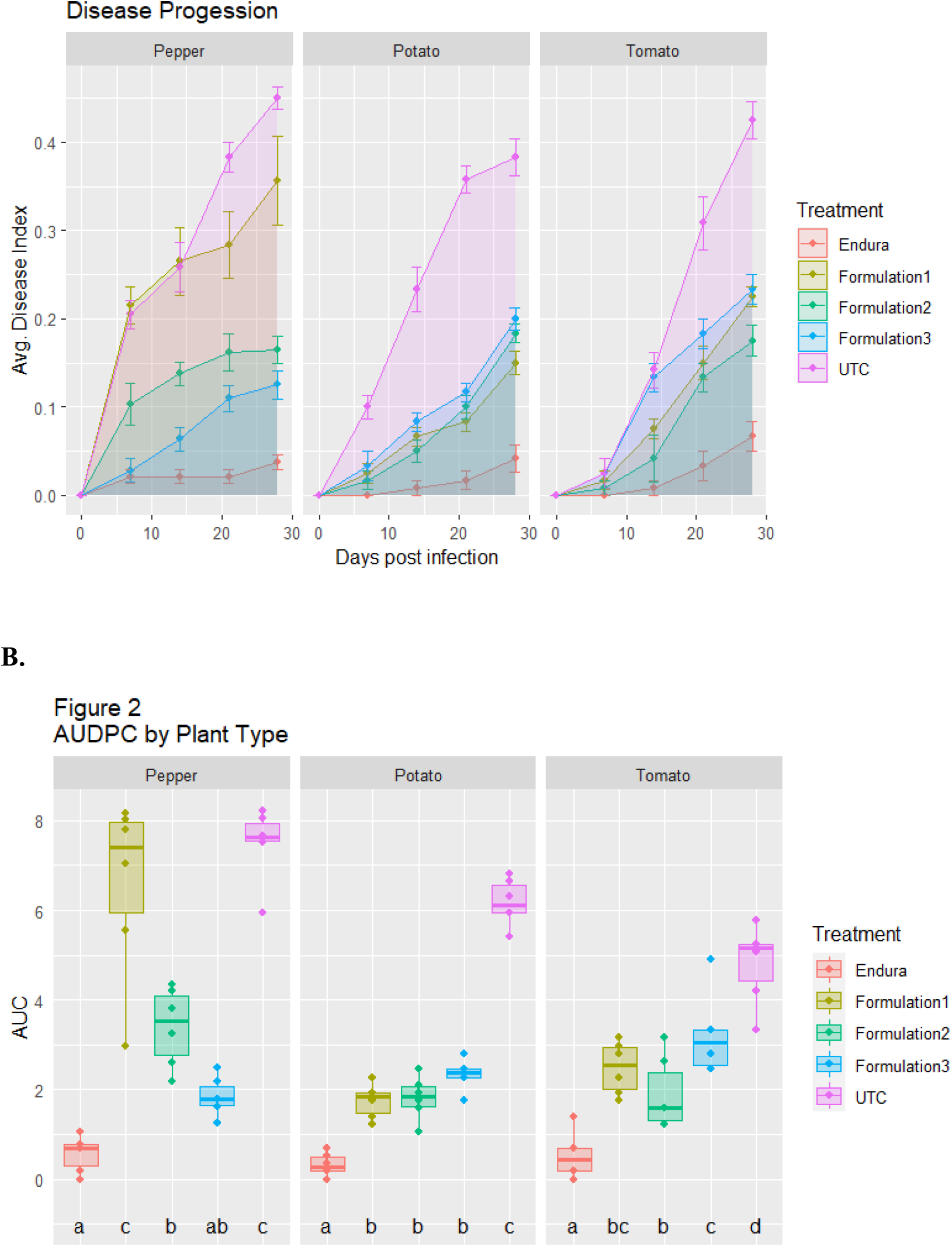
Analysis of the area under the disease progress curve. A) The disease progression is displayed with error bars calculated using the standard deviation, plotted using the disease severity. B) An analysis of variance (ANOVA) test was conducted with 95% confidence intervals. The disease severity is plotted, with the symbols representing induvial treatments used.

### 2.5 Experimental design

A total of 90 plants were used for this investigation: 30 tomato, 30 potato, and 30 bell pepper plants. For each crop there were 6 replications done for 5 treatments (Table 1). The plants were examined for previous disease, physical damage, or pest infestations in the laboratory. All plants were 15 cm in height and were approximately 10 weeks old at the start of the investigation. The plants were placed in separate greenhouses based on the type of plant at 25 °C and were placed in 1-gallon pots with Speedling^®^ potting mix and with 1 cup of 2 tablespoons of N-P-K fertilizer diluted with 1-gallon of water.

### 2.6 Application of the Treatments

In this study, the preventative efficacy of the treatments was tested, so the treatments were sprayed 24 hours before the inoculum was sprayed. On each plant the treatments and the early blight inoculum were sprayed until run-off using a hand-held sprayer.

The industrial standard fungicides used in this investigation were Endura^®^, to treat the early blight fungus in the bell pepper crop and Inspire Super^®^ to treat the early blight fungus in the tomato and potato crops.

### 2.7 Measuring the Disease Severity

The disease severity was measured weekly for 28 days and was measured by a proportion of total foliage infected to the total foliage (Figure 2). Infected foliage was classified as a leaf showing any symptoms of early blight: lesions, black or brownish concentric rings, and the chlorosis of the leaf, to note the most common symptoms of early blight.

### 2.8 Statistical Analysis

An analysis of variance (ANOVA) test, with a confidence interval of 95%, and an alpha value of 0.05. The null hypothesis (H0), p-value is greater than 0.05, states that there is no statistically significant difference between the AUDPC of the control and treated crops. The alternate hypothesis (H_A_), if p-value is less than 0.05, states that there is a statistically significant difference between the control crops and the treated crops.

### 2.9 Estimating the Area Under the Disease Progress Curve

The area under the disease progress curve (AUDPC) was calculated for each treatment using the trapezoidal rule. The area under the disease progress curve is used by plant pathologists to determine the progression of a disease over a certain period as a reference for disease resistance. The AUDPC can be found by taking the integral of the disease progress curve, however the most customary practice is to use the formula in Figure 5. The AUDPC formula uses the trapezoidal method of calculating the area under the curve. The trapezoidal method involves taking the average of the disease severity between 2 adjacent data points and multiplying the result by the time interval and continuing this process for each set of time intervals until the total area is determined (Madden et al., 2007).

**Figure 5.**
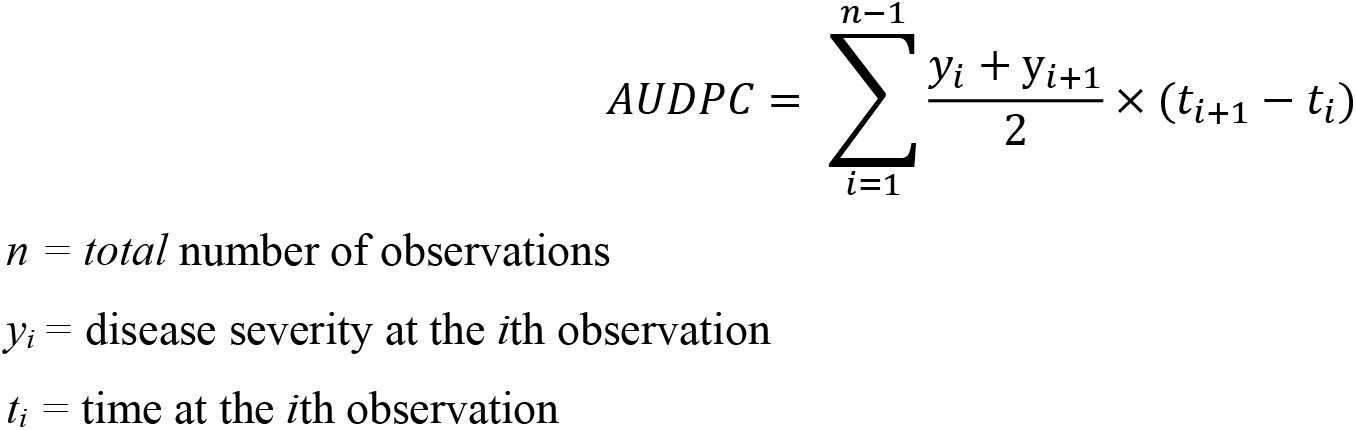
AUDPC formula.

## 3. Results and Discussion

**Table 5.** The AUDPC values

| **Treatments** | **AUDPC** |
| --- | --- |
| Formulation 1 | 247.91 |
| Formulation 2 | 189.58 |
| Formulation 3 | 320.83 |
| Industry Standard | 52.5 |
| Untreated Control | 481.25 |

**B.**
| **Treatments** | **AUDPC** |
| --- | --- |
| Formulation 1 | 175 |
| Formulation 2 | 122.5 |
| Formulation 3 | 110.83 |
| Industry Standard | 32.08 |
| Untreated Control | 627.08 |

| **Treatments** | **AUDPC** |
| --- | --- |
| Formulation 1 | 659.16 |
| Formulation 2 | 339.5 |
| Formulation 3 | 82.25 |
| Industry Standard | 56.875 |
| Untreated Control | 822.5 |

### 3.1 Data and Statistics

The AUDPC, is a measure for the disease severity over a period of time. Low AUDPC values over the period of 28 days suggested that the microbial treatments were effective in treating early blight, reflected in the disease progression graphs. As expected, the industry standard, had the least disease progression value in all three crops, with an AUDPC value of 32.08 in potato, 52.5 in tomato, and 10 in bell pepper; however, the trends displayed in the disease progression showed that in the potato crop, formulation 1 had effectively controlled the disease, with an AUDPC value of 175. In the tomato crop, formulation 2 had the least disease progression, with an AUDPC value of 189.58. In the bell pepper crop, formulation 3 performed the most effectively against early blight with an average AUDPC value of 115.17.

## 4. Conclusion

### 4.1 Ecological Application and Human Health

These Endura^®^ and Inspire Super^®^ are too toxic to use regularly and risk the potential development of resistance against the fungicides. The EPA labels for these fungicides contain a surface water advisory, limiting the application of these fungicides near shallow water tables or lakes due to the potential of run-off accumulation after months of application (Kish, 2008; Garvie, 2019).

The microbial formulations can be sprayed in rotation with the industry-standard fungicides to reduce the risk of developing pathogenic resistance and reduce the chemical run-off of fungicidal treatments on the surrounding water bodies (Zubrod et al., 2019; Hahn, 2014). Additionally, only using the microbial formulations would eliminate the chemical run-off.

The chemicals used in common fungicides are also known to cause health issues through exposure to skin, ingestion, and inhalation (N.A. Smart, 2003).

In this study Endura^®^ was used as the industrial standard to treat early blight in the bell pepper crop. Endura^®^ contains the active ingredient boscalid, which has low acute toxicity through the oral, dermal, and inhalation routes but, has the liver and thyroid as the target organs in mice and dogs (Giles-Parker, 2013). In mice, exposure to boscalid resulted in fatty changes in the liver and in dogs it resulted in an increase in alkaline phosphatase levels and in hepatic weights (Giles-Parker, 2013). The EPA also suggests that boscalid is suggestive of having evidence of carcinogenicity (Giles-Parker, 2013).

Inspire Super^®^ was another industrial standard used to treat early blight in the tomato and potato crops. Inspire Super^®^ contains the active ingredient difenoconazole which also has a low acute toxicity through the oral, dermal, and inhalation routes but, is a mild eye irritant (PMRA, 2015). When animals such as mice and rats were given repeated doses, it was reported that there were effects on the liver, body weight, and food consumption of the animals (PMRA, 2015). It was also noted that when mice were given excessive doses of difenoconazole the formation of liver tumors were observed, however the same effect was not present in mice (PMRA, 2015). When difenoconazole was given to pregnant animals, serious effects were noted: increase incidence in fetal mortality and mothers had depressed body weight gains (PMRA, 2015).

### 4.2 Bacillus subtilis 29784, Bacillus subtilis 168

In the field of plant pathology, the *Bacillus subtilis* species is one of the most studied microorganisms, as it has the potential to be a single mode of action fungicide (Li and Leifert, 1994). The *Bacillus subtilis* species produces lipopeptides which are amphiphilic secondary metabolites classified in 3 different families: fengycin, iturin, and surfactin (Hoffmann et al., 2021) Out of these three families, surfactin is synthesized by the *Bacillus subtilis 168* and *29784* microorganisms as an immune response from fungal infections, and has strong antifungal activities (Krishnan et al., 2019). The exact mode of action of surfactin is unknown, however, this lipopeptide is known to trigger immune responses in the host tissues (Henry et al., 2011). Both *Bacillus subtilis 29784* and *Bacillus subtilis 168* are part of the surfactin family, identified by aβ-hydroxyl fatty acid (C_12_–C_16_) (Ntushelo et al., 2019). Current investigations have examined the effects of the surfactin against *Fusarium moniliforme* and grape downy mildew, however, have not been tested on early blight, which is the purpose of selecting *Bacillus subtilis 168* and *Bacillus subtilis 29784* for this investigation (Li et al., 2019; Jiang et al., 2016).

### 4.3 Azotobacter sp

The genus *Azotobacter* is a common microorganism found in neutral to alkaline soils and has been widely used as a biofertilizer (Gerlach and Vogel, 1902). *Azotobacter sp.* are gram negative bacteria that are known for the symbiotic biological nitrogen fixation process and are used as biological fertilizers to address agricultural challenges such as nutrient deficiencies in the soil (Sumbul et al., 2020). The process of biological nitrogen fixation allows the conversion of atmospheric nitrogen (N_2_) into ammonium which can be absorbed by the roots, which improves the structural and physiological composition of the plant (Aasfar et al., 2021). *Azotobacter sp.* can contribute to efficient crop production, especially in developing countries where there is a limited supply of mineral fertilizers, can be used in N-fertilizers to combat this issue, or can potentially be used in organic pesticides or fungicides to treat and stimulate growth in the plant (Sumbul et al., 2020). When *Azotobacter sp.* is placed in high-stress environmental conditions, it results in cyst formation to combat environmental stress, which makes the microorganism beneficial to use in extreme environmental conditions (Aasfar et al., 2021). Using *Azotobacter sp.* in microbial formulation 1 would incorporate the multiple modes of action, because of the combination of the antifungal properties of *Bacillus subtilis 29784* and *Bacillus subtilis 168* and the biological nitrogen fixation properties of *Azotobacter sp.* (Hu et al., 2013).

### 4.4 *Glomus claroideum*, *Glomus etunicatum*, and *Glomus viscosum*

The *Glomus* genus of fungi are classified as effective microbes: effective microbes help increase crop growth through increasing plant productivity and photosynthetic efficiency (Birhane et al., 2012). Current studies have focused on the effects of effective microbes on sunflowers where it has been reported that effective microbes increase chlorophyll, N, P, carbohydrate, and protein contents (Balliu et al., 2015). The *Glomus* genus of fungi are more specifically classified as arbuscular mycorrhiza fungi (AMF) which form a symbiotic relationship with the roots: the symbiosis between the microorganism and the roots allows for an increase in photosynthetic rate and an increase in water uptake (Walker and Vestberg, 1998). AMF function by changing the morpho-physiological traits through forming a hyphal network with the plant increasing root surface area and improving plant health and growth (Begum et al., 2019). AMF is commonly used in microbial formulations as a biofertilizer to increase plant health in areas with high abiotic stress and used for sustainable agricultural practices (Kumar et al., 2022). The *Glomus* genus was used in this study because of the stimulation of plant growth it provides, which results in a multiple modes of action response when these microorganisms are paired with antifungal microorganisms (Hu et al., 2013).

### 4.5 Trichoderma viride

*Trichoderma viride* has been tested to show antifungal and biocontrol responses on many common agricultural diseases such as *Phytophthora, Sclerotinia,* and *Fusarium* (Zin and Badaluddin, 2020). The genus *Trichoderma* produces hydrolytic enzymes and secondary metabolites which helps to treat the spores of plant parasitic fungi (Zin and Badaluddin, 2020). *Trichoderma viride* controls the development of many fungal diseases by producing antibiotics, volatile compounds, and inducing plant prime resistance (Alfiky and Weisskopf, 2021). Biological mechanisms are also used by *Trichoderma viride* to control plant parasitic fungi: competition for nutrients, ecological niche, and antibiosis (Zin and Badaluddin, 2020). *Trichoderma viride* has multiple modes of action because although the most common use of this microorganism is as a fungicide, *Trichoderma viride* also solubilizes phosphates in the soil through producing enzymes (Kubheka and Ziena, 2021). In a study analyzing the effects of adding *Trichoderma viride* on sugar cane, the nitrogen, potassium, phosphorus and organic carbon intake increased after the inoculation with *Trichoderma viride,* indicating that *Trichoderma viride* has the potential to increase nutrient uptake while exhibiting antifungal properties (Sood et al., 2020) however, *Trichoderma viride* would still need an additional microorganism to be paired with to significantly increase immune system responses and plant growth (Yadav et al., 2009).

### 4.6 Acetobacter sp

*Acetobacter sp.* is commonly used to prevent sour rot, a disease formed on injured berries, primarily grapes (Ivey et al., 2021). The *Acetobacter genus.* is commonly used as a plant biofertilizer because of the ability of the microorganism to fix nitrogen and promote the expansion of the root system (Sumbul et al., 2020). The *Acetobacter* produces Indole Acetic Acid (IAA) and Gibberellic acid (GA) which causes the increase in the number of rootlets and root proliferation (Egamberdieva et al., 2017). An additional mode of action of *Acetobacter* is phosphate solubilization, which increases phosphorus absorption in agro-ecosystems and further develops the metabolic pathways in plants (Sashidhar and Podile, 2010). *Acetobacter sp.* forms symbiotic relationships with plants and expands the root surface area increasing nutrient absorption (de Souza et al., 2015). Currently, there is a commercially available biofertilizer Katyayni^®^, however, *Acetobacter sp.* has not been tested or commercially developed in combination with antifungal microorganisms.

### 4.7 Future Applications and Research

In the future, the three formulations will be field-tested over a longer time and contain more replications to determine the effectiveness of the formulations in the field. If the microbial formulations show more promising results in the field, these formulations can be submitted to the EPA for further evaluation and use in the field. Additionally, the mode of action of the formulation as well as of the secondary metabolites of these microorganisms, would need to be studied to observe the interactions of the microorganisms when in a formulation, currently research on the mode of action of single microorganism has been done, but the mode of action has not been observed when the microorganisms are in a formulation, which could lead to an explanation of the varying levels of effectiveness of each microbial formulation based on the plant type.

